# Spatial and compositional variations in fruit characteristics of papaya (*Carica papaya* cv. Tainung No. 2) during ripening

**DOI:** 10.1101/2023.01.31.526316

**Authors:** Sun Woo Chung, Seolah Kim, Seong Cheol Kim

## Abstract

Papaya fruit (*Carica papaya*) has different degrees of ripening within a fruit, affecting its commercial market value. The fruit characteristics of ‘Tainung No. 2’ papaya was investigated at the stem-end, middle, and calyx-end parts at three ripening stages and categorized based on fruit skin coloration: unripe at ca. 16 weeks after anthesis (WAA), half-ripe at ca. 18 WAA, and full-ripe at ca. 20 WAA. The fruits maintained an elliptical shape during ripening with 2.36 of the ratios of the length to the width. The peel and pulp color changed from green to white to yellow during ripening, regardless of the fruit three parts. In the pulp, soluble solid content increased to about 320% and firmness decreased to about 99% during ripening but did not differ among fruit three parts. Individual nutrient contents, including primary and secondary metabolites, and minerals, changed dynamically between the ripening stages and fruit parts. Total carbohydrates and proteins, N, and K, were more accumulated at the stem-end during ripening, meanwhile fructose, glucose, Mg, and Mn were at the calyx-end. In the principal component analysis, ripening stages and fruit parts were distinctly determined by the first and second principal components, respectively. These results provide fundamental information for improving ripening during papaya cultivation.

## 1. Introduction

Papaya (*Carica papaya*) is a perennial herbaceous crop widely cultivated for its sweet and flavored fruit. The global production of papaya fruit reached 14.10 million metric tons in 2021, making it the fourth most popular tropical fruit behind banana, mango, and pineapple (FAO, 2023). Papaya originated from the lowland of Mexico to Panama and is found primarily in tropical regions between 23 °N and S latitudes because the cultivation requires high temperatures year-round (Daagema *et al*., 2020). Subtropical regions, including Mediterranean countries, can also cultivate papaya; some of these regions have to use protected facilities, including a greenhouse because low temperatures during winter affect fruit set, growth, and production (Gunes and Gübbük, 2012; Salinas *et al*., 2019; Salinas *et al*.,2022). Recently, papaya cultivation has expanded from subtropical and tropical regions to temperate regions, including Japan and the Republic of Korea, owing to increasing average temperature and consumer needs (Jeon *et al*., 2022; Jeong *et al*., 2020; Ogata *et al*., 2016).

Papaya fruit quality is determined by various characteristics required from the market. Papaya fruits have different shapes as sex forms, including males, females, and hermaphrodites. Markets prefer the hermaphrodite fruit shape, which tends to be pear-shaped or elongated (Daagema et al., 2020). The nutritional value of papaya fruit depends on the fruit ripening stages. Papaya fruit ripening is accompanied by physiochemical changes, including sugar metabolism, peel color changes, and pulp softening (Farina *et al*., 2020). The more ripened fruit showed different physicochemical characteristics, such as moisture, titratable acidity, and total soluble solids, as well as higher antioxidant activity and higher total phenolic and flavonoid contents (Jeon et al., 2022; Zuhair *et al*., 2013). Minerals according to fruit ripening progress have different distributions depending on the fruit ripening stage (Chukwuka *et al*., 2013). The fruit is consumed regardless of ripeness and can be either raw or in processed forms (Dotto and Abihudi, 2021) with unripe fruits being used as a vegetable and ripe fruits and ripe fruit (Gunes and Gübbük, 2012). However, the ripening degree within a fruit would be different from the stem end to the calyx end due to subtle differences, including the degrees of sun exposure and fruit skin temperatures, which has been observed in blueberries (Spinardi *et al*., 2019), apples (Doerflinger *et al*., 2015; Singha *et al*., 1991), and grapes (Ryu *et al*., 2020). The different ripening degrees of fruit parts may lead to a loss of consumer confidence. However, the changes in papaya fruit have not been well studied compared to those in other fruits.

In this study, we investigated morphological, physicochemical, and nutritional characteristics in different parts, ranging from stem end to calyx end, of ‘Tainung No.2’ papaya fruits during ripening. We also investigated the relationship between the fruit characteristics, parts, and ripening stages. The fruit characteristics and their relationships would be contributed to a commercially valuable database associated with papaya fruit physiology.

## 2. Materials and methods

### 2.1. Plant materials

‘Tainung No. 2’ papaya seeds were sowed on November 10th, 2015, in trays at a greenhouse of an experimental orchard in the Research Institute of Climate Change and Agriculture, the National Institute of Horticultural and Herbal Science, Jeju, Republic of Korea (33° 28 N’, 126° 31’ E) and then transplanted on June 8th, 2016, into the ground of the same greenhouse. These trees were cultivated according to the standard guidelines for papaya cultivation (Joa *et al*., 2012): the minimum temperature during winter was maintained at 15 °C in the greenhouse by using hot air blowers to prevent papaya trees from chilling injury (Fig. 1).

**Fig. 1.**
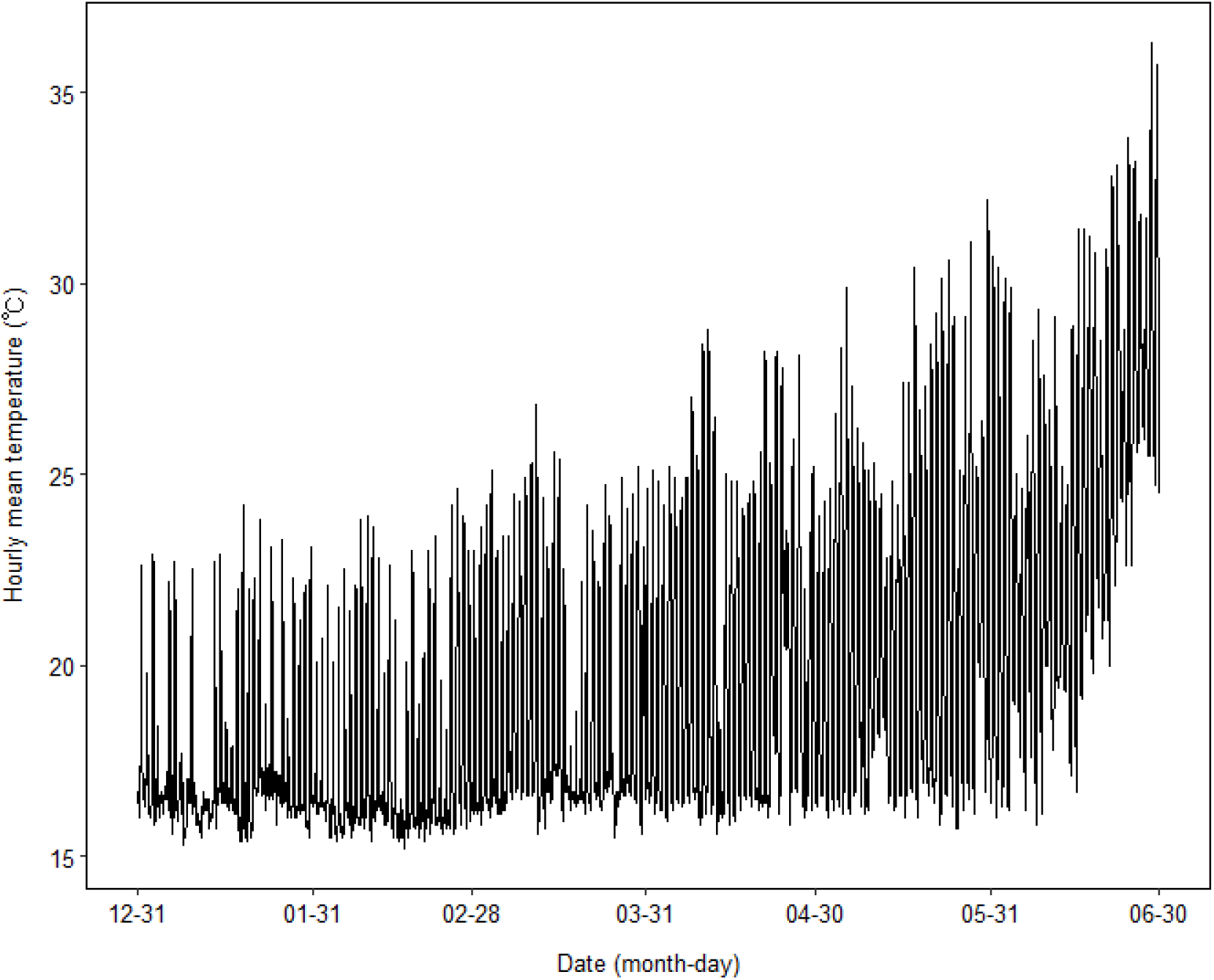
Changes in temperature and relative humidity in the greenhouse for papaya (*Carica papaya*) cultivation

The fruit was categorized into three ripening stages based on weeks after anthesis (WAA), according to Zuhair et al. (2013): (1) unripe at ca. 16 WAA, (2) half-ripe at ca. 18 WAA, and (3) full-ripe at ca. 20 WAA. At each stage, three fruits per a tree were harvested, which was performed at three trees to provide three biological replicates. After the size and weight of each fruit were measured, each fruit was dissected into three parts of equal lengths: stem-end, middle, and calyx-end for further analyses.

### 2.2. Determination of colors, soluble solid contents (SSC), and firmness

The peel and pulp colors of papaya fruits were measured at the stem end, middle, and calyx end at the three ripening stages using a spectrophotometer (CM700d, Minolta Co., Osaka, Japan) and described by the CIE L*, a*, and b* color space coordinates (McGuire, 1992). The L* value represents the lightness of colors, ranging from 0 to 100 (0, black; 100, white). The a* value was negative for green and positive for red. The b* value was negative for blue and positive for yellow. The values were measured at three midpoint regions of each fruit part. In addition to the CIE coordinates, The hue angle (h°) was calculated as tan^-1^ (b*/a*), which implies visual color appearance; 0°, red-purple; 90°, yellow; 180°, bluish-green; 270°, blue. The chroma (C*) was computed as (a*^2^ + b*^2^)^1/2^, which describes the quality of the color intensity or saturation.

The SSC was measured using a digital refractometer (HI98801, Hanna Instruments Inc., Woonsocket, RI, USA). Peel firmness was measured using a texture analyzer (TA-XT express, Stable Micro System, UK). A puncture test was conducted using a 2.0 mm diameter cylindrical probe (Stable Micro System) at three different points along the fruit equator with a cross-head speed of 2.0 mm s^-1^ and a penetration depth of 5.0 mm.

### 2.3. Determination of water proportion and carbohydrate, protein, and lipid contents

Fruits were dehydrated at 105 °C in a drying oven up to constant weight for determining the proportion of water content compared with the total fresh weight of each sample, which was expressed as a percentage (%). To determine carbohydrate, protein, and lipid contents, all the samples were lyophilized using a freeze dryer and were finely ground using a mortar and pestle. Carbohydrate was determined according to the procedure of Jermyn (1975). Protein concentration was determined using a Micro-Kjeldahl’s apparatus (Chromý *et al*., 2015). Lipid concentration was determined using the Soxhlet extraction method (Luque de Castro and Priego-Capote, 2010). All analyses were performed according to the official analysis methods of the Association of Official Analytical Chemists (AOAC) (Cunniff and Washington, 1997).

### 2.4. Quantification of free sugars

Free sugars was extracted according to the method described by Oh *et al*. (2011) with some modifications. Three grams of ground fruits were added to 50 mL of 50% acetonitrile. The samples were extracted thrice by ultrasonic extraction method for 6 h. The extract was diluted and cleaned up using a Sep-Pak C18 cartridge column (Waters, Milford, MA, USA) and filtered through a 0.45 μm pore size micro filter (Woongki Science Co., Ltd., Seoul, Korea). After the sample preparation, free sugars were separated in Prevail TM Carbohydrate ES column (4.6 × 250 mm, 5 μm, Grace, Deerfield, IL, USA) equipped with an HPLC system (Waters 2695, Waters Associate Inc., Milford, MA, USA). The eluents were passed through the column at a flow rate of 0.8 mL/min using acetonitrile: distilled water (70:30, v/v). The chromatographic peak corresponding to free sugar was identified by comparing the retention times with those of glucose, fructose, and sucrose (Sigma, St. Louis, MO, USA) as standards. Concentrations were calculated using the calibration curve generated from the standard solutions.

### 2.5. Quantification of ascorbic acids

Ascorbic acid was extracted using the procedure described by Rizzolo *et al*. (1984) with some modifications. Thirty grams of samples were added to 25 mL of 6% metaphosphoric acid. The sample was homogenized and then centrifuged at 10,000 × *g* at 4°C for 10 min. The supernatant was filtrated through a filter paper (Whatman No. 4, Whatman plc, Kent, UK) and a membrane filter (MF-millipore^TM^ 0.22 μm, Merck KGaA, Darmstadt, Germany). The solution was cleaned up through a Waters Sep-Pak C18 column (Waters, Milford, CT, USA). Ascorbic acids in the prepared samples were separated using a Waters Bonda Pak NH2 (3.9 × 300 mm) column (Waters) equipped with an HPLC system (Waters TM 600 controller, Waters TM 616 Pump, Waters 717 plus auto Sampler, Waters TM 486 tunable absorbance detector). The eluent was passed through the column at a flow rate of 1 mL/min using 5 mM KH2PO4 (pH 4.6): acetonitrile (30:70, v/v). The chromatographic peak corresponding to ascorbic acid in the samples was identified by comparing its retention times with those of the standard (Sigma). Concentrations were calculated using the calibration curve generated from standard solutions prepared for the standard.

### 2.6. Determination of macro and micro elements

The proportions of nitrogen and phosphorus contents were determined using the Kjeldahl method (Bradstreet, 1954) and the molybdenum blue method (Crouch and Malmstadt, 1967), respectively. Three grams of fruits were weighed and heated in a furnace for 4 h at 550°C. The crucible was cooled down in a desiccator and dissolved in 2.5 mL of a decomposition solution (HNO_3_: H_2_SO_4_: HClO_4_ (10:1:4, v/v/v)). The solution was filtered and diluted up to 100 mL using distilled water. Exchangeable cations (K, Ca, and Mg) and minerals (Fe, Mn, Zn, and Cu) were determined by inductively coupled plasma spectrophotometry (ICP-Integra XL, GBC Scientific Equipment Pty Ltd, Australia) following the procedure of Zarcinas *et al*.(1987). The results were obtained by using a working standard of 1,000 ppm for each sample (Oh et al., 2011).

### 2.7. Statistical analyses

Statistical analyses were performed using the R 4.2.2 (R Core Team, 2022). Statistical significances were determined using ANOVA with the agricolae v1.3-5 package (de Mendiburu, 2021). Means were compared using Tukey HSD tests at *p* < 0.5.

Principal components analysis was carried out on samples by replicate means to obtain a correlation among fruit characteristics using the factoextra v1.0.7 package (Kassambara and Mundt, 2020). The variables were standardized by subtracting the mean and then dividing by the standard deviation of each original variable to assign each weight in the analysis.

## 3. Results

The fruit length, width, the ratio of the length to the width, and weight did not change during ripening (Table 1). The length of papaya fruit was an average of 23.6 mm ranging from 19.2 to 25.8 mm; the width was 10.0 mm ranging from 9.6 to 10.3 mm. The ratio of the length to width ranged from 1.9 to 2.7, indicating the elliptical shape of the papaya fruit. The fresh weight of the fruit ranged from 699.6 to 997.9 g with an average of 854.5 g.

**Table 1.** Fruit length, width, their ratio, and weight of papaya (*Carica papaya*) during ripening.

| Stage | Length | Width | Ratio of length to width | Weight |
| --- | --- | --- | --- | --- |
|  | mm |  |  | g |
| Unripe | $25.9 \pm 1.40$ a<sup>z</sup> | $9.6 \pm 0.54$ a | $2.7 \pm 0.27$ a | $865.9 \pm 123.92$ a |
| Half-ripe | $25.8 \pm 0.80$ a | $10.3 \pm 1.11$ a | $2.5 \pm 0.32$ a | $997.9 \pm 110.92$ a |
| Full-ripe | $19.2 \pm 5.08$ a | $10.0 \pm 0.73$ a | $1.9 \pm 0.67$ a | $699.6 \pm 200.26$ a |

<sup>z</sup>Different letters within columns indicate significant differences by Duncan's multiple range test. **Fig. 2.** Inner and outer of papaya (*Carica papaya*) fruit in different ripening parts: Unripe (left), half-ripe (middle), full-ripe (right).**Table 2**Peel chromaticity of papaya (*Carica papaya*) in three parts during ripening.

The peel color of the papaya fruit changed during ripening (Figure 2 and Table 2). The peel lightness decreased at the half-ripe and full-ripe stages, compared with the unripe stage (Table 2). The half-ripe and full-ripe stages become redder during ripening compared to the unripe stage. The b value increased during ripening, indicating that the pulp becomes yellower. The h° between the parts did not change at each ripening stage. The h° decreased in the half-ripe and full-ripe stages, compared to that in the unripe stage; the values were the average of 129.9, 75.5, and 58.0 at the unripe, half-ripe, and full-ripe stages, respectively. The h° of the unripe stage was significantly higher than those of the other two stages. The h° tended to decrease during fruit ripening continuously. The trend was obviously shown in the calyx-end parts of each stage, where the h° significantly decreased. The C* increased in the half-ripe and full-ripe stages (51.9 and 60.0, respectively), compared to the unripe stage (13.2); there was no significant difference between the half-ripe and full-ripe stages.

**Fig. 2.**
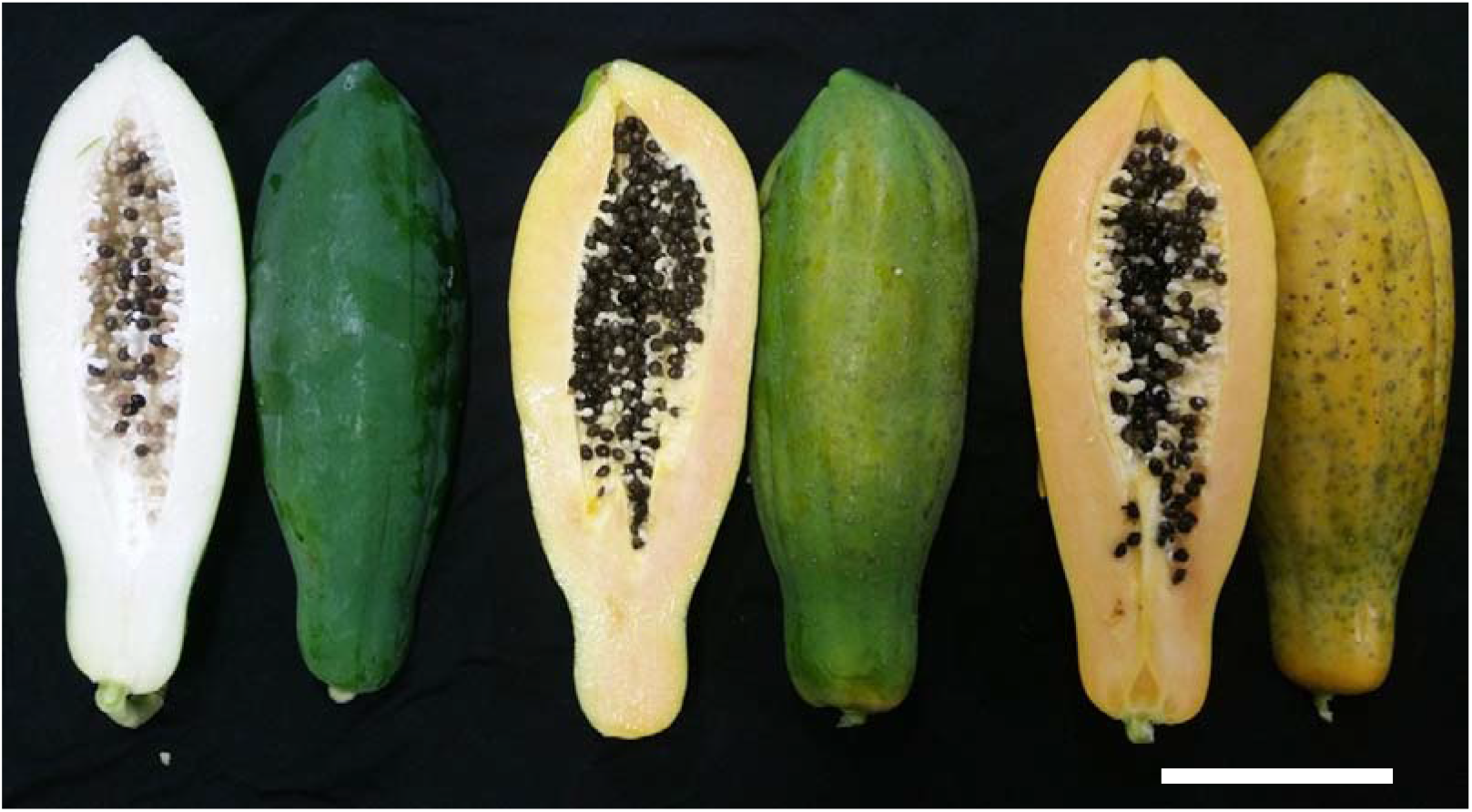
Inner and outer of papaya (*Carica papaya*) fruit in different ripening parts: Unripe (left), half-ripe (middle), full-ripe (right).

**Table 2.**
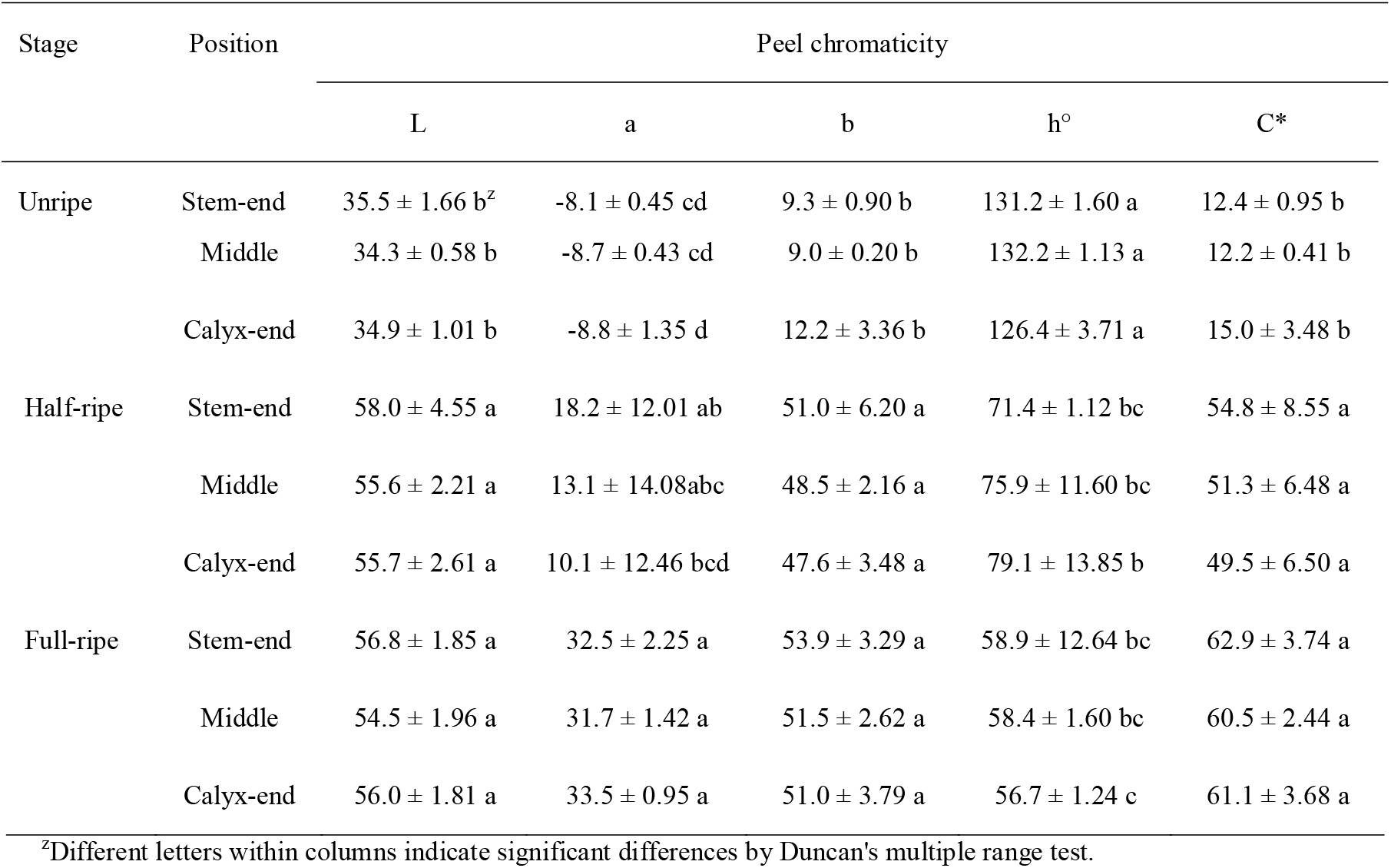
Peel chromaticity of papaya (*Carica papaya*) in three parts during ripening.

Pulp characteristics were investigated during fruit parts and ripening (Table 3). The pulp color changed from unripe to half-ripe stages (Table 3), but not afterward. The lightness became darker in the half-ripe and full-ripe stages (55.8) than in the unripe stages (75.1). The a and b values of the unripe stage were lower than those of the half-ripe and full-ripe stages, indicating greenish and yellowish color, respectively. The significance of the h° and C* during ripening was consistent with the L, a, and b values. SSC did not change between the fruit parts at each ripening stage but changed significantly during ripening, from the unripe stage (3.9 °Brix) to the half-ripe and full-ripe stages (10.8 °Brix). Pulp firmness did not change at the fruit parts in each ripening stage. The firmness did not change between the unripe and half-ripe stages and decreased only at the full-ripe stage up to 96% (from 9.75 N to 0.37 N).

**Table 3.**
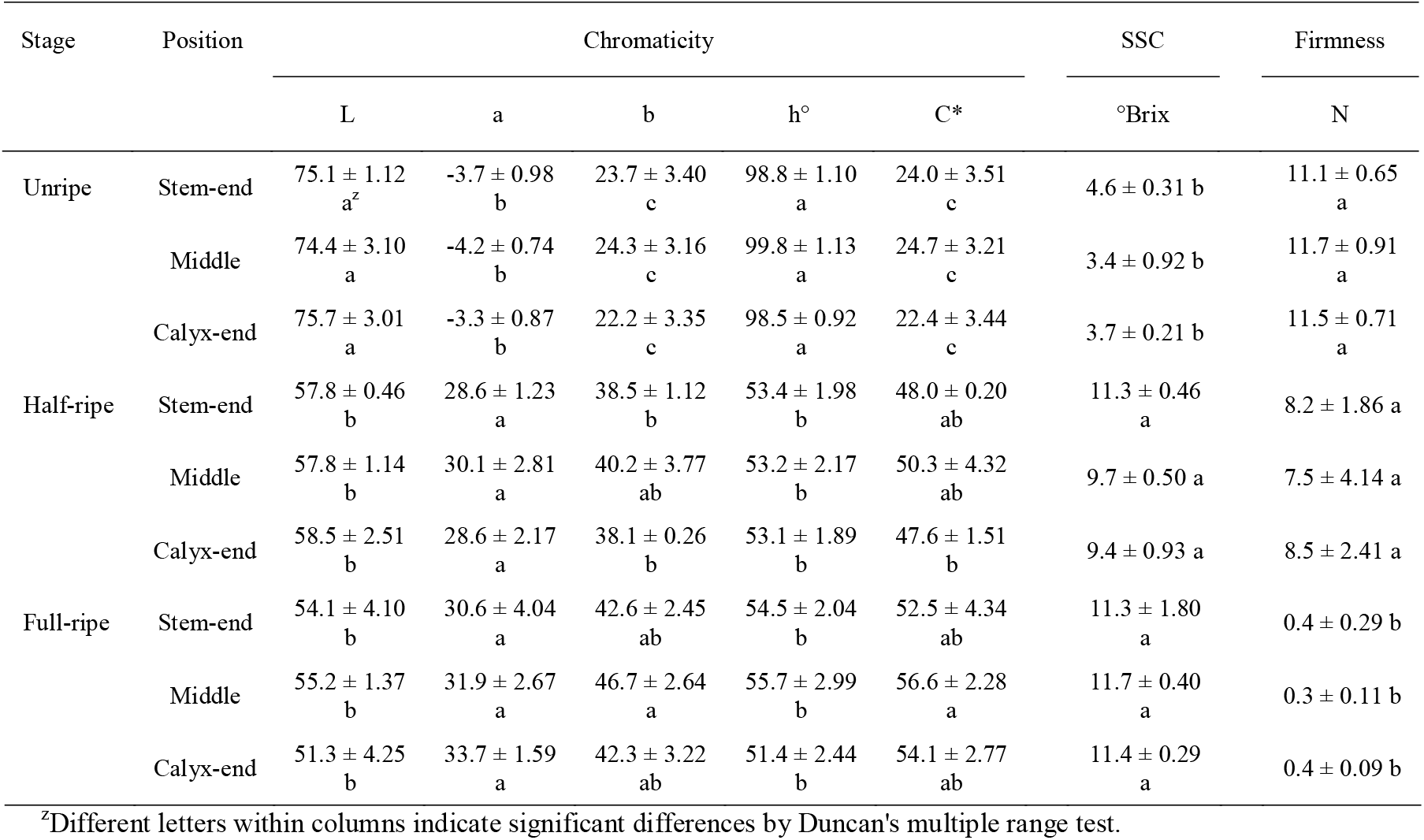
Pulp soluble solid content (SSC), firmness, and chromaticity of papaya (*Carica papaya*) at three parts during ripening.

| Stage | Position | Chromaticity |  |  |  |  | SSC | Firmness |
| --- | --- | --- | --- | --- | --- | --- | --- | --- |
|  |  | L | a | b | h° | C* | °Brix | N |
| Unripe | Stem-end | 75.1 ± 1.12<br>a<sup>z</sup> | -3.7 ± 0.98<br>b | 23.7 ± 3.40<br>c | 98.8 ± 1.10<br>a | 24.0 ± 3.51<br>c | 4.6 ± 0.31 b | 11.1 ± 0.65<br>a |
|  | Middle | 74.4 ± 3.10<br>a | -4.2 ± 0.74<br>b | 24.3 ± 3.16<br>c | 99.8 ± 1.13<br>a | 24.7 ± 3.21<br>c | 3.4 ± 0.92 b | 11.7 ± 0.91<br>a |
|  | Calyx-end | 75.7 ± 3.01<br>a | -3.3 ± 0.87<br>b | 22.2 ± 3.35<br>c | 98.5 ± 0.92<br>a | 22.4 ± 3.44<br>c | 3.7 ± 0.21 b | 11.5 ± 0.71<br>a |
| Half-ripe | Stem-end | 57.8 ± 0.46<br>b | 28.6 ± 1.23<br>a | 38.5 ± 1.12<br>b | 53.4 ± 1.98<br>b | 48.0 ± 0.20<br>ab | 11.3 ± 0.46<br>a | 8.2 ± 1.86 a |
|  | Middle | 57.8 ± 1.14<br>b | 30.1 ± 2.81<br>a | 40.2 ± 3.77<br>ab | 53.2 ± 2.17<br>b | 50.3 ± 4.32<br>ab | 9.7 ± 0.50 a | 7.5 ± 4.14 a |
|  | Calyx-end | 58.5 ± 2.51<br>b | 28.6 ± 2.17<br>a | 38.1 ± 0.26<br>b | 53.1 ± 1.89<br>b | 47.6 ± 1.51<br>b | 9.4 ± 0.93 a | 8.5 ± 2.41 a |
| Full-ripe | Stem-end | 54.1 ± 4.10<br>b | 30.6 ± 4.04<br>a | 42.6 ± 2.45<br>ab | 54.5 ± 2.04<br>b | 52.5 ± 4.34<br>ab | 11.3 ± 1.80<br>a | 0.4 ± 0.29 b |
|  | Middle | 55.2 ± 1.37<br>b | 31.9 ± 2.67<br>a | 46.7 ± 2.64<br>a | 55.7 ± 2.99<br>b | 56.6 ± 2.28<br>a | 11.7 ± 0.40<br>a | 0.3 ± 0.11 b |
|  | Calyx-end | 51.3 ± 4.25<br>b | 33.7 ± 1.59<br>a | 42.3 ± 3.22<br>ab | 51.4 ± 2.44<br>b | 54.1 ± 2.77<br>ab | 11.4 ± 0.29<br>a | 0.4 ± 0.09 b |

<sup>z</sup>Different letters within columns indicate significant differences by Duncan's multiple range test.

The water contents continuously decreased during fruit ripening (Table 4). The water contents of unripe, half-ripe, and full-ripe stages were 93.9, 92.5, and 91.2%, respectively. The water content was not different among the parts of the unripe and half-ripe stages. In the full-ripe stage, the content was the lowest in the stem end (90.6%) over the other parts (91.5%). The total carbohydrates tended to increase during fruit ripening, showing the differences among the parts (Table 4). The lowest amount was 4.5 g/100g at the stem-end and the calyx-end of the unripe stage; conversely, the highest was 7.4 g/100g at the stem-end of the full-ripe stage. The accumulation rate of carbohydrates at the stem end increased 120% to 137% during ripening, while the middle ranged from 120% to 110%, and at the calyx end ranged from 131% to 117%. Protein contents did not show distinct tendencies during ripening in part of the fruit, except that the accumulation from the stem-end of each stage was higher than that in the other parts. In addition, the lipid contents were not detected in any fruit part during the ripening.

**Table 4.** Water, carbohydrate, protein, and lipid contents of papaya (*Carica papaya*) at three parts during ripening.

| Stage | Position | Water | Carbohydrate | Protein | Lipid |
| --- | --- | --- | --- | --- | --- |

|  |  | % | g/100g |  |  |
| --- | --- | --- | --- | --- | --- |
| Unripe | Stem-end | 94.0 ± 0.03 a<sup>z</sup> | 4.5 ± 0.02 e | 1.06 ± 0.03 cd | nd<sup>y</sup> |
|  | Middle | 93.7 ± 0.15 a | 5.1 ± 0.15 d | 0.91 ± 0.01 de | nd |
|  | Calyx-end | 93.9 ± 0.02 a | 4.5 ± 0.05 e | 1.15 ± 0.05 bc | nd |
| Half-ripe | Stem-end | 92.6 ± 0.02 b | 5.4 ± 0.05 d | 1.29 ± 0.01 ab | nd |
|  | Middle | 92.5 ± 0.04 b | 6.1 ± 0.02 c | 0.93 ± 0.03 de | nd |
|  | Calyx-end | 92.3 ± 0.03 b | 5.9 ± 0.03 c | 1.11 ± 0.01 c | nd |
| Full-ripe | Stem-end | 90.6 ± 0.08 d | 7.4 ± 0.08 a | 1.39 ± 0.02 a | nd |
|  | Middle | 91.4 ± 0.04 c | 6.7 ± 0.03 b | 1.20 ± 0.02 bc | nd |
|  | Calyx-end | 91.6 ± 0.05 c | 6.9 ± 0.04 b | 0.88 ± 0.01 e | nd |

<sup>z</sup>Different letters within columns indicate significant differences by Duncan's multiple range test.
<sup>y</sup>Nd means not detected.

Three soluble sugars were determined, and two soluble sugars were detected during ripening (Table 5). Sugar contents increased during ripening and were different from that of the fruits. Fructose contents were not different among the parts at the unripe stage: the fructose contents in the calyx end of the two other stages were approximately 110% higher than that in the middle and stem end. The glucose content in each part of the fruit significantly increased during ripening. However, these contents were not different at the three parts within each ripening stage, except for the full-ripe stage, where the glucose content of the calyx end were higher than that of the stem end.

**Table 5.** Changes in the content of soluble sugars at each part of papaya pulp during ripening.

| Stage | Position | Fructose | Glucose | Sucrose |
| --- | --- | --- | --- | --- |
|  |  | g·100 g<sup>-1</sup> |  |  |
| Unripe | Stem-end | 1.1 ± 0.01 e<sup>z</sup> | 1.3 ± 0.00 d | nd<sup>y</sup> |
|  | Middle | 1.1 ± 0.08 e | 1.3 ± 0.03 d | nd |
|  | Calyx-end | 1.4 ± 0.07 e | 1.6 ± 0.03 d | nd |
| Half-ripe | Stem-end | 1.8 ± 0.17 d | 2.0 ± 0.07 c | nd |
|  | Middle | 1.9 ± 0.23 d | 2.1 ± 0.09 c | nd |
|  | Calyx-end | 2.1 ± 0.12 c | 2.4 ± 0.04 c | nd |
| Full-ripe | Stem-end | 2.5 ± 0.01 bc | 2.9 ± 0.01 b | nd |
|  | Middle | 2.7 ± 0.14 ab | 3.1 ± 0.17 ab | nd |
|  | Calyx-end | 2.9 ± 0.05 a | 3.4 ± 0.05 a | nd |

<sup>z</sup>Different letters within columns indicate significant differences by Duncan's multiple range test.<sup>y</sup>Nd means not detected.**Table 6**

In addition to these structural compounds, bioactive compounds and minerals are essential components of fruits. In the papaya fruit, ascorbic acid contents increased up to approximately 300 % from unripe to half-ripe and approximately 140 % from half-ripe to full-ripe in all three parts during ripening (Table 6). However, there were no differences between the ripening stages. The proportion of mineral compounds varied between fruit parts and ripening stages (Table 7 and 8). The K proportion was about 2.85% during ripening, followed by N (0.80%), P (0.37%), Ca (0.24%), Mg (0.13%), Fe (26.05 ppm), Cu (2.73 ppm), Zn (11.11 ppm), and Mn (1.13 ppm). The proportion of K in each part was maintained during ripening, whereas those of other minerals were fluctuated during ripening. The change patterns of N proportion were distinctly different for each part: increase at the stem-end in the full-ripe stage, decrease at the middle in the half-ripe stage, decrease and then increase at the calyx-end in the half-ripe and full-ripe stage, respectively. The P proportion changed only at the stem-end between the unripe and full-ripe stages. The Ca proportion steadily decreased at the stem-end and middle parts during ripening; that of the calyx-end decreased in the half-ripe stage and was maintained up to the full-ripe stage. The Mg proportion of all fruit parts decreased from the unripe to half-ripe stages and then remained in the full-ripe stage. The Fe proportion of all the parts decreased from the unripe to half-ripe stages. Subsequently, the Fe proportion of stem-end did not change and those of the middle and calyx end increased at the full-ripe stage. The Cu proportion decreased only in the calyx end from the unripe to half-ripe stages; all the other proportions did not differ between the parts and ripening stages. The Zn proportion decreased from the unripe to the half-ripe stage and was maintained between the parts and ripening stages. The Mn proportion at the stem-end did not change during ripening, that at the middle increased only in the full-ripe stage, and that at the calyx-end decreased and then increased during ripening.

**Table 6.** Changes in the content of ascorbic acid at each part of papaya pulp during ripening.

| Stage | Position | Ascorbic acid |
| --- | --- | --- |
|  |  | mg·L<sup>-1</sup> |
| Unripe | Stem-end | 8.7 ± 0.43 c<sup>z</sup> |
|  | Middle | 6.8 ± 0.20 c |
|  | Calyx-end | 6.9 ± 0.37 c |
| Half-ripe | Stem-end | 21.9 ± 0.23 b |
|  | Middle | 23.2 ± 0.53 b |
|  | Calyx-end | 22.1 ± 1.10 b |
| Full-ripe | Stem-end | 32.7 ± 0.97 a |
|  | Middle | 31.4 ± 0.97 a |
|  | Calyx-end | 30.7 ± 1.13 a |

<sup>z</sup>Different letters within columns indicate significant differences by Duncan's multiple range test.**Table 7**

**Table 7.** Change of macronutrients from fruit part in ripening part of papaya fruit: N, nitrogen; P, phosphorous; K, potassium; Ca, Calcium; Mg, magnesium.

| Stage | Position | N | P | K | Ca | Mg |
| --- | --- | --- | --- | --- | --- | --- |

|  |  | % |  |  |  |  |
| --- | --- | --- | --- | --- | --- | --- |
| Unripe | Stem-end | $0.7 \pm 0.01$ def<sup>z</sup> | $0.5 \pm 0.02$ a | $3.3 \pm 0.13$ a | $0.4 \pm 0.03$ a | $0.15 \pm 0.010$ b |
| | Middle | $0.8 \pm 0.01$ bcd | $0.4 \pm 0.02$ bc | $2.8 \pm 0.10$ b | $0.4 \pm 0.04$ a | $0.20 \pm 0.010$ a |
| | Calyx-end | $1.2 \pm 0.07$ a | $0.4 \pm 0.03$ b | $2.6 \pm 0.13$ b | $0.3 \pm 0.02$ b | $0.16 \pm 0.010$ b |
| Half-ripe | Stem-end | $0.7 \pm 0.08$ ef | $0.4 \pm 0.01$ ab | $3.5 \pm 0.19$ a | $0.3 \pm 0.02$ bc | $0.09 \pm 0.000$ e |
| | Middle | $0.6 \pm 0.02$ f | $0.3 \pm 0.01$ c | $2.7 \pm 0.14$ b | $0.3 \pm 0.02$ bc | $0.12 \pm 0.000$ cd |
| | Calyx-end | $0.8 \pm 0.04$ cde | $0.4 \pm 0.02$ b | $2.6 \pm 0.10$ b | $0.2 \pm 0.01$ cd | $0.13 \pm 0.000$ c |
| Full-ripe | Stem-end | $0.9 \pm 0.07$ bc | $0.4 \pm 0.04$ bc | $3.3 \pm 0.25$ a | $0.2 \pm 0.01$ d | $0.09 \pm 0.000$ e |
| | Middle | $0.7 \pm 0.01$ def | $0.3 \pm 0.03$ c | $2.5 \pm 0.19$ b | $0.1 \pm 0.00$ d | $0.11 \pm 0.000$ de |
| | Calyx-end | $0.9 \pm 0.07$ b | $0.4 \pm 0.02$ ab | $2.5 \pm 0.14$ b | $0.1 \pm 0.00$ d | $0.13 \pm 0.000$ c |

<sup>z</sup>Different letters within columns indicate significant differences by Duncan's multiple range test.

**Table 8.**
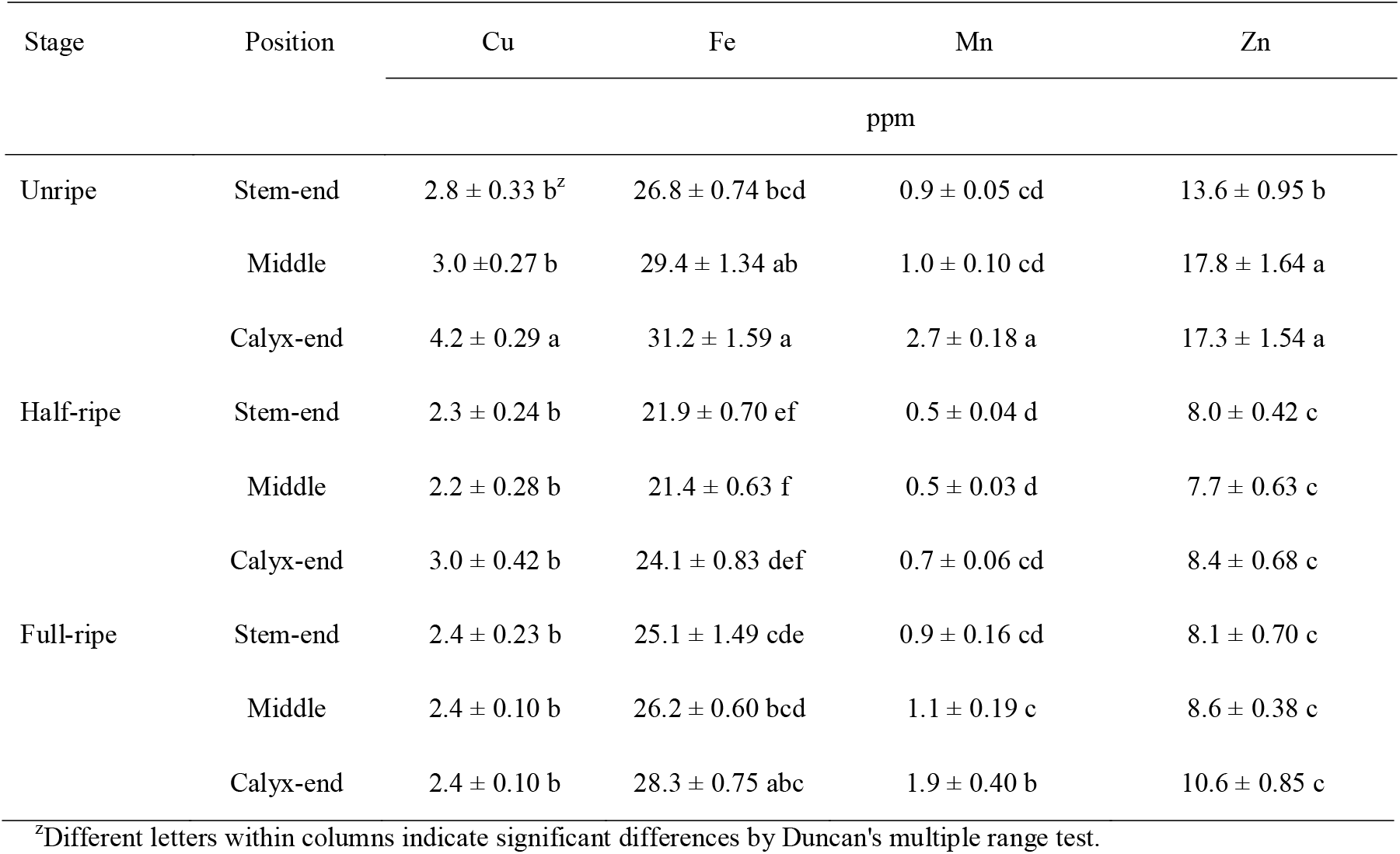
Change of micronutrients from ripening part of papaya fruit.

The fruit characteristics of papaya were located in each principal component (PC), which was arranged according to the size of variance (Table 9, Fig 3, and Fig 4). The five principal components explained 98.33% of all the variances, and PC1 and PC2 accounted for 85.36% (Table 9 and Fig. 3). PC1 explained 71.78% with the eigenvalue of 19.38 (Table 9 and Fig. 3) and represented the chromaticity, SSC, and ascorbic acid. These fruit characteristics were approximately 0.98 of the correlation value on absolute average and contributed to 59.78% in PC1. Of these, peel h^o^, pulp L, and pulp h^o^ were negatively correlated with PC1, whereas the others were positively correlated.

**Fig 3.**
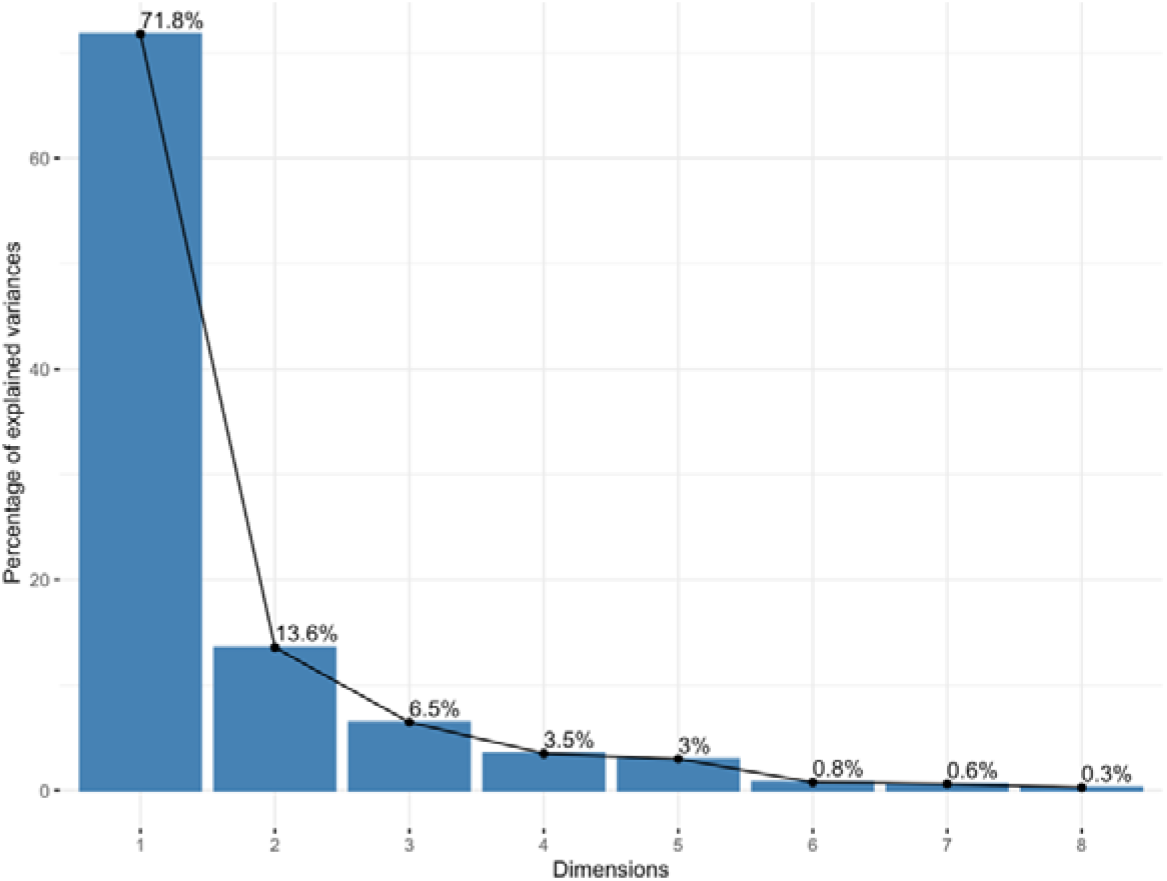
Scree plot for the percentage of explained variance of principal components.

**Fig. 4.**
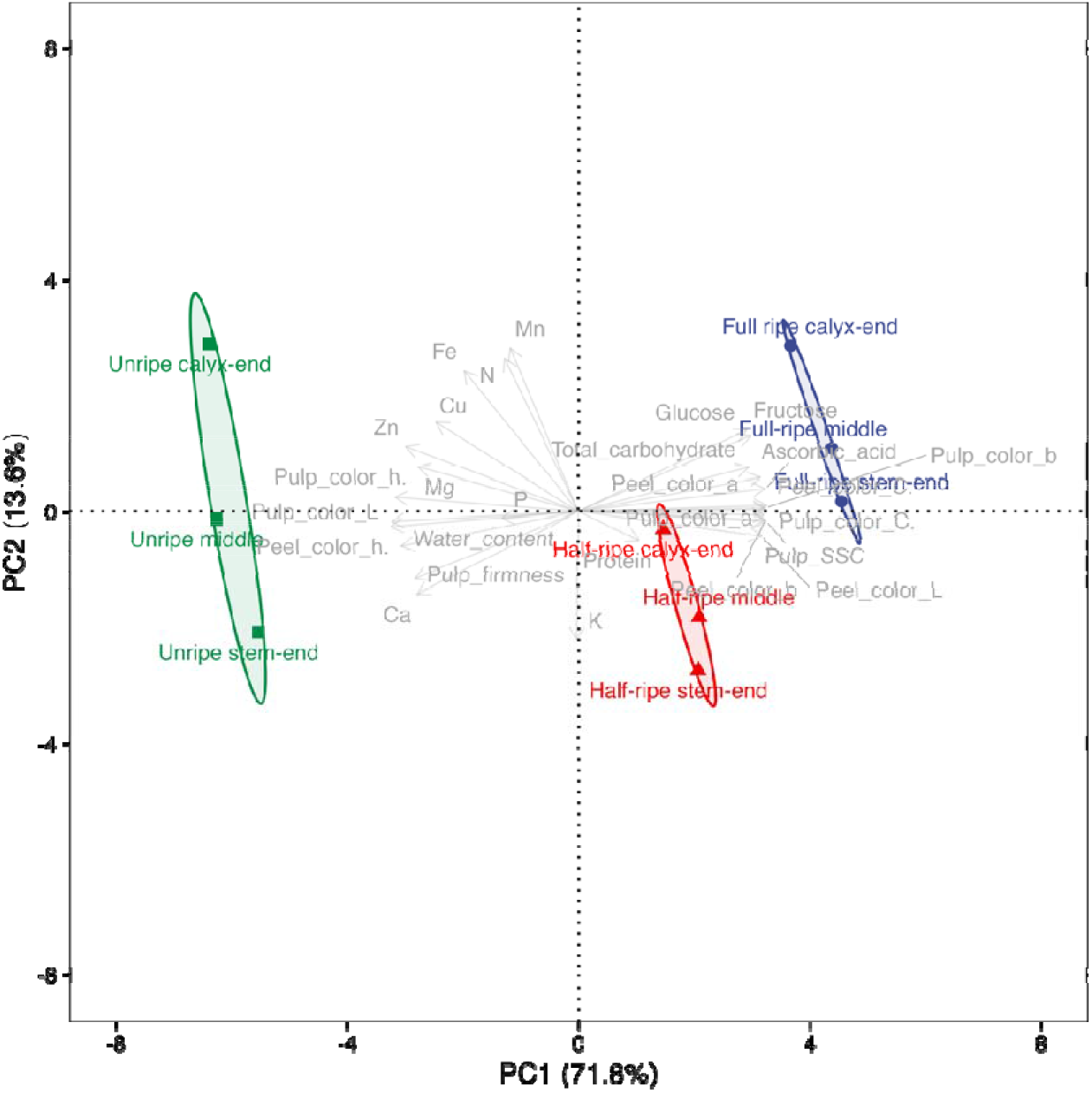
Plot of fruit qualities and part of principal component (PC) scores of means on the first two PC axes for the fruit part of papaya (*Carica papaya*) during ripening with 95% confidence of ellipses for the mean of ripening stages. Percentages in parenthesis of each axis represent variances of each PC.

**Table 9.**
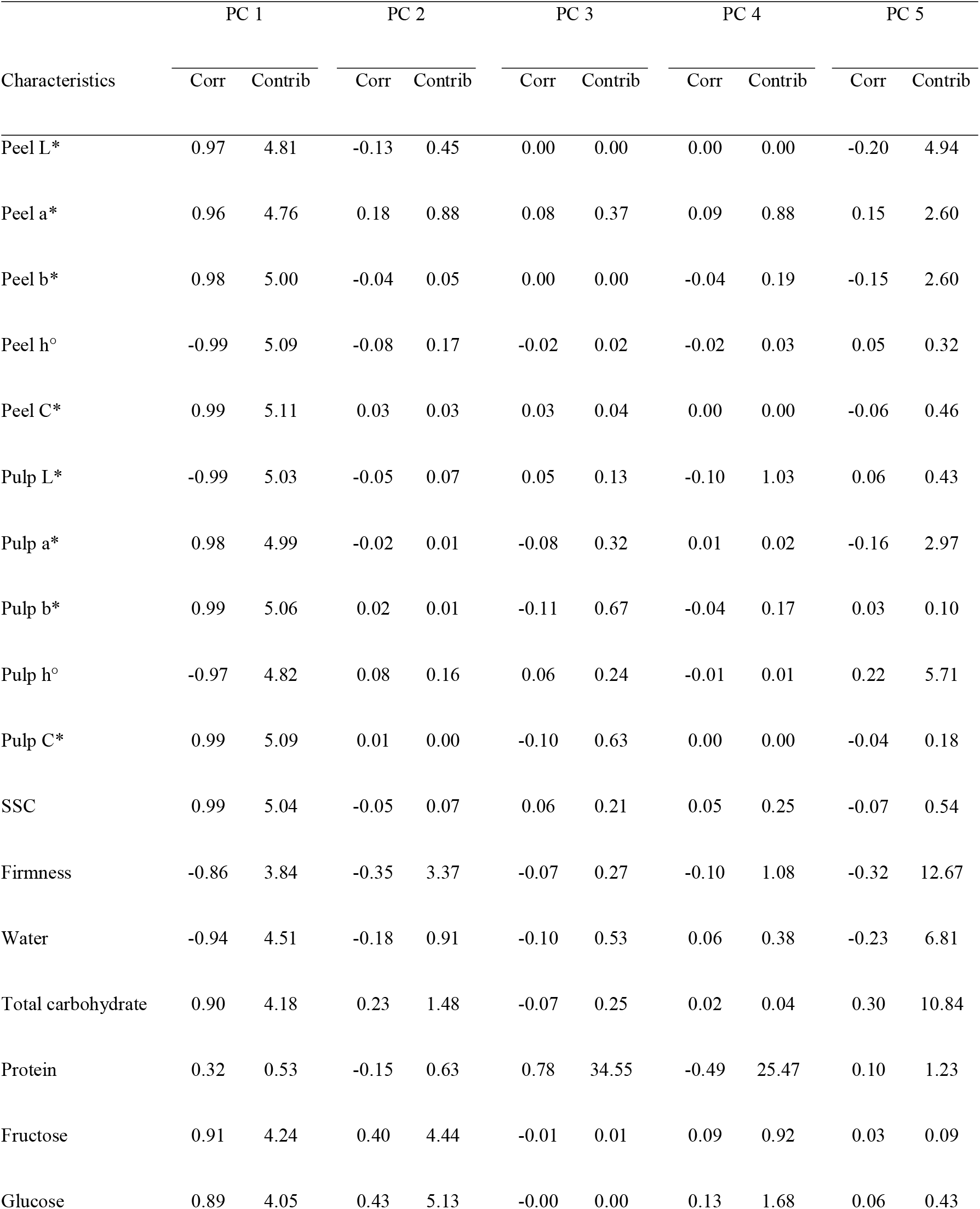
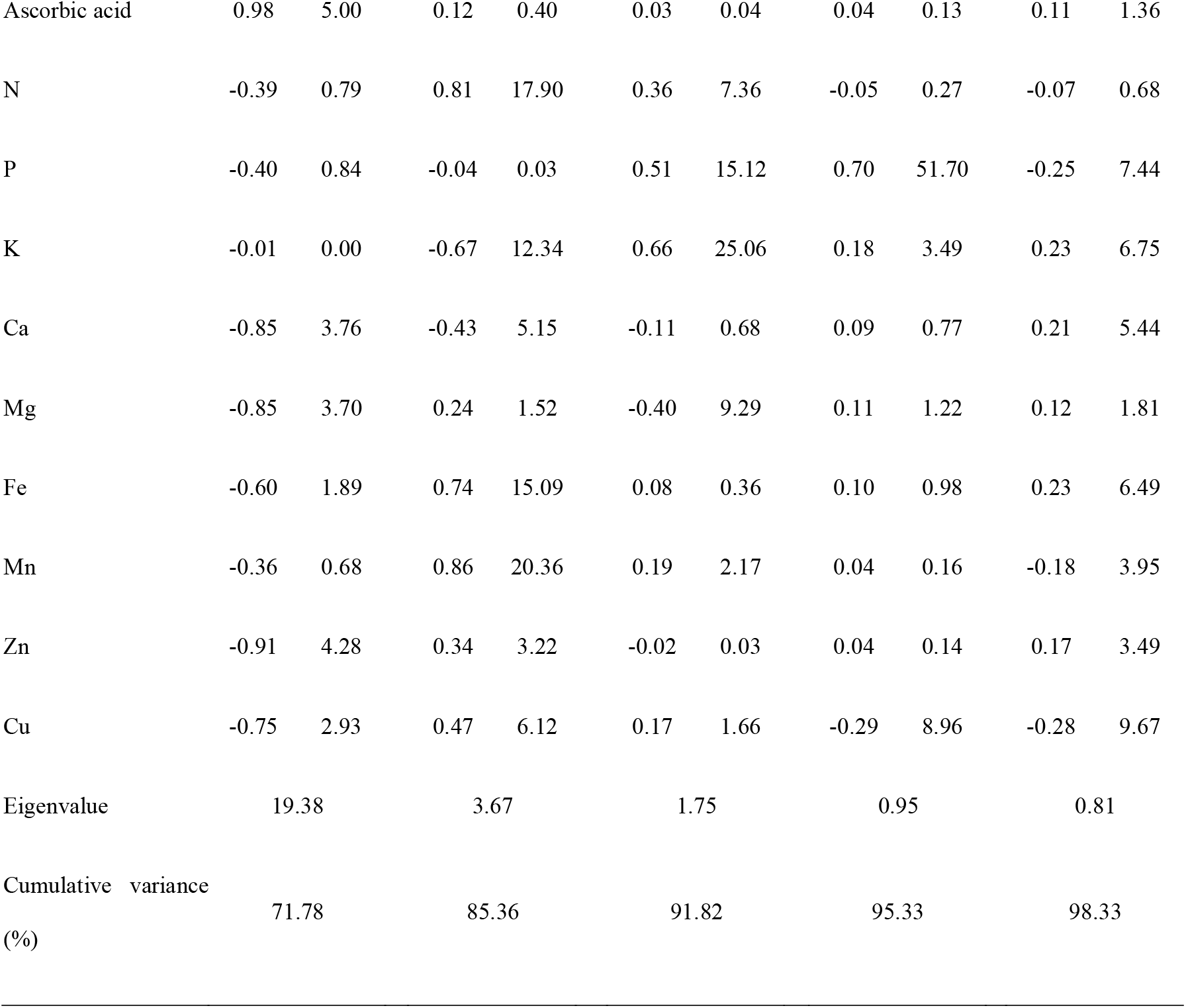
Correlation (Corr) and contribution (Contrib) values of fruit characteristics of papaya (*Carica papaya*) at each principal component (PC) from PC analysis of the fruit part during ripening. The eigenvalues and cumulative percentage of variation are listed at the bottom of the columns

|  | PC 1 |  | PC 2 |  | PC 3 |  | PC 4 |  | PC 5 |  |
| --- | --- | --- | --- | --- | --- | --- | --- | --- | --- | --- |
| Characteristics | Corr | Contrib | Corr | Contrib | Corr | Contrib | Corr | Contrib | Corr | Contrib |
| Peel L* | 0.97 | 4.81 | -0.13 | 0.45 | 0.00 | 0.00 | 0.00 | 0.00 | -0.20 | 4.94 |
| Peel a* | 0.96 | 4.76 | 0.18 | 0.88 | 0.08 | 0.37 | 0.09 | 0.88 | 0.15 | 2.60 |
| Peel b* | 0.98 | 5.00 | -0.04 | 0.05 | 0.00 | 0.00 | -0.04 | 0.19 | -0.15 | 2.60 |
| Peel h° | -0.99 | 5.09 | -0.08 | 0.17 | -0.02 | 0.02 | -0.02 | 0.03 | 0.05 | 0.32 |
| Peel C* | 0.99 | 5.11 | 0.03 | 0.03 | 0.03 | 0.04 | 0.00 | 0.00 | -0.06 | 0.46 |
| Pulp L* | -0.99 | 5.03 | -0.05 | 0.07 | 0.05 | 0.13 | -0.10 | 1.03 | 0.06 | 0.43 |
| Pulp a* | 0.98 | 4.99 | -0.02 | 0.01 | -0.08 | 0.32 | 0.01 | 0.02 | -0.16 | 2.97 |
| Pulp b* | 0.99 | 5.06 | 0.02 | 0.01 | -0.11 | 0.67 | -0.04 | 0.17 | 0.03 | 0.10 |
| Pulp h° | -0.97 | 4.82 | 0.08 | 0.16 | 0.06 | 0.24 | -0.01 | 0.01 | 0.22 | 5.71 |
| Pulp C* | 0.99 | 5.09 | 0.01 | 0.00 | -0.10 | 0.63 | 0.00 | 0.00 | -0.04 | 0.18 |
| SSC | 0.99 | 5.04 | -0.05 | 0.07 | 0.06 | 0.21 | 0.05 | 0.25 | -0.07 | 0.54 |
| Firmness | -0.86 | 3.84 | -0.35 | 3.37 | -0.07 | 0.27 | -0.10 | 1.08 | -0.32 | 12.67 |
| Water | -0.94 | 4.51 | -0.18 | 0.91 | -0.10 | 0.53 | 0.06 | 0.38 | -0.23 | 6.81 |
| Total carbohydrate | 0.90 | 4.18 | 0.23 | 1.48 | -0.07 | 0.25 | 0.02 | 0.04 | 0.30 | 10.84 |
| Protein | 0.32 | 0.53 | -0.15 | 0.63 | 0.78 | 34.55 | -0.49 | 25.47 | 0.10 | 1.23 |
| Fructose | 0.91 | 4.24 | 0.40 | 4.44 | -0.01 | 0.01 | 0.09 | 0.92 | 0.03 | 0.09 |
| Glucose | 0.89 | 4.05 | 0.43 | 5.13 | -0.00 | 0.00 | 0.13 | 1.68 | 0.06 | 0.43 |
| Ascorbic acid | 0.98 | 5.00 | 0.12 | 0.40 | 0.03 | 0.04 | 0.04 | 0.13 | 0.11 | 1.36 |
| N | -0.39 | 0.79 | 0.81 | 17.90 | 0.36 | 7.36 | -0.05 | 0.27 | -0.07 | 0.68 |
| P | -0.40 | 0.84 | -0.04 | 0.03 | 0.51 | 15.12 | 0.70 | 51.70 | -0.25 | 7.44 |
| K | -0.01 | 0.00 | -0.67 | 12.34 | 0.66 | 25.06 | 0.18 | 3.49 | 0.23 | 6.75 |
| Ca | -0.85 | 3.76 | -0.43 | 5.15 | -0.11 | 0.68 | 0.09 | 0.77 | 0.21 | 5.44 |
| Mg | -0.85 | 3.70 | 0.24 | 1.52 | -0.40 | 9.29 | 0.11 | 1.22 | 0.12 | 1.81 |
| Fe | -0.60 | 1.89 | 0.74 | 15.09 | 0.08 | 0.36 | 0.10 | 0.98 | 0.23 | 6.49 |
| Mn | -0.36 | 0.68 | 0.86 | 20.36 | 0.19 | 2.17 | 0.04 | 0.16 | -0.18 | 3.95 |
| Zn | -0.91 | 4.28 | 0.34 | 3.22 | -0.02 | 0.03 | 0.04 | 0.14 | 0.17 | 3.49 |
| Cu | -0.75 | 2.93 | 0.47 | 6.12 | 0.17 | 1.66 | -0.29 | 8.96 | -0.28 | 9.67 |
| Eigenvalue | 19.38 |  | 3.67 |  | 1.75 |  | 0.95 |  | 0.81 |  |
| Cumulative variance<br>(%) | 71.78 |  | 85.36 |  | 91.82 |  | 95.33 |  | 98.33 |  |

## 4. Discussion

Papaya fruits have different shapes as sex forms, including male, female, and hermaphrodite. Markets prefer the hermaphrodite fruit shape, which tends to be pear-shaped or elongated (Araya-Valverde *et al*., 2019; Salinas et al., 2019). In our result, the ratio of length to width of ‘Tainung No.2’ was an average of 2.4 during the entire ripening stage (Table 1), which is consistent with the variety’s general shape as hermaphroditic type (Kung *et al*., 2010). In addition to this ratio, the variables related to fruit volume did not change during ripening (Table 1). The size development of papaya fruit is completed before the initiation of ripening (Salinas et al., 2019). A sigmoidal curve was observed in papaya ripening patterns in various papaya varieties (Salinas et al., 2019). This would be associated with the suspension of cell expansion and enlargement when using carbon sources for metabolite changes, including pigment and soluble sugar accumulations (Doerflinger et al., 2015; Seymour *et al*., 2013).

The degree of fruit ripeness of fruits is a determining factor in consumption, marketing, and processing (Barragán-Iglesias *et al*., 2018). Fruit color is an indicator of the degree of ripening degree in fresh fruits, including papaya (Barragán-Iglesias et al., 2018), blueberries (Chung *et al*., 2016; Chung *et al*., 2019), grapes (Ryu et al., 2020), and mangoes (Chung *et al*., 2021), which also provides visual cues for marketable values. ‘Tainung No.2’ papaya fruit showed distinctly different peel and pulp colors (Fig. 2). The colors were evenly distributed throughout the whole fruit at each ripening stage (Tables 2 and 3). In addition to morphological characteristics, SSC increased, and firmness decreased during ripening, regardless of the fruit parts (Table 2). These changes in SSC and firmness are commonly observed during the ripening of various fruits, including papaya (Barragán-Iglesias et al., 2018; Gomez *et al*., 2002) and mangoes (Chung et al., 2021). In papaya fruit, the pulp softening is highly correlated with the sweetness process associated with SSC, probably because of the easier release of cellular contents in the fully ripened tissue (Gomez et al., 2002).

In the present study, individual metabolites in papaya fruit showed various accumulation patterns in each fruit part during ripening. Research on the differences among different tissue zones has mainly been conducted on apples (Angmo *et al*., 2022; Doerflinger et al., 2015; Lee *et al*., 2019; Lewis, 1980). To the best of our knowledge, this is the first report to determine the characteristics of each part of the papaya fruit; there are no comparative data in previous studies on papaya fruits. Doerflinger et al. (2015) compared the carbohydrate concentration of the stem end, middle, and calyx end in three apple cultivars (‘Empire’, ‘Honeycrisp’, and ‘Gala’) during ripening. In ‘Empire’, the concentration of starch was highest at the calyx end and lowest at the stem end. ‘Honeycrisp’ and ‘Empire’ had the highest concentration of sorbitol in the calyx end, whereas it was highest in the stem end in ‘Gala’. The distribution differences of glucose, fructose, and sucrose were similar in all three cultivars: higher fructose and glucose concentrations in the stem end and higher sucrose concentrations in the calyx end of the fruit. In papaya fruit, soluble sugars were accumulated mainly when the fruit remains attached to the plant (Gomez et al., 2002). At the unripe stage, glucose is prevalent among the soluble sugars, and during ripening, sucrose becomes the predominant sugar with the modification of the soluble sugar profiles (Paull, 1996). Chan Jr and Kwok (1975) reported that sucrose levels varied from 1.8% to 8.0% during ripening. In this study, ‘Tainung No. 2’ papaya fruit accumulated more total carbohydrates in the stem end than in the other parts (Table 4). The main soluble sugars in papaya are glucose, fructose, and glucose (Kelebek *et al*., 2015); however, their compositions vary among cultivars (Kelebek et al., 2015). In ‘Tainung No. 2’ papaya fruit, only fructose and glucose were identified during ripening, and more accumulated in the calyx end (Table 5). In addition to carbohydrates, more protein was accumulated at the stem end. These differences could be associated with different development rates in different tissue zones. An increase in total primary amounts is associated with carbon allocation from the photosynthetic organs before ripening. Meanwhile, the accumulation of soluble sugars is one of the ripening processes. In papaya fruit, the accumulation rate rapidly increases during ripening (Selvaraj *et al*., 1982). Therefore, the ripening process is more activated regarding changes in primary metabolites. The differences in the structure and size of cells could also lead to developmental differences within the fruit. However, such differences vary by species (Doerflinger et al., 2015) and its cultivars (Bain and Robertson, 1951; Leshem *et al*., 1984).

Establishing nutrient absorption and accumulation patterns is critical for planning optimum nutrient supply and improving the influence on fruit quality (Casero *et al*., 2017). Macro and micro element accumulation show dynamic variance during ripening. In this study, all elements except N at the stem end decreased or maintained at the full-ripe stage, despite different patterns at the half-ripe stage among fruit parts. The decreases of the elements had also been shown in various fruits, including papaya (Chukwuka et al., 2013) and apple fruit (Mota *et al*., 2022; Nachtigall and Dechen, 2006). In apple pulp, when comparing fruits approximately 60 and 120 days after full bloom for the elements N, P, K, Mg, and S, there is a slight decrease, but Ca, Fe, Cu, Mn, and B contents had no significant differences (Mota et al., 2022). In papaya pulp, Ca, K, P, and Mg decreased to 75.56%, 38.67%, 66.46%, and 50.00%, respectively, at the full-ripe stage, compared to the unripe stage. (Chukwuka et al., 2013).

All papaya fruit samples were distinctly divided according to fruit characteristics (Fig. 4). While ripening stages were accounted for by PC1, including peel and pulp colors, fruit parts were accounted for by PC2. Even though the fruit characteristics were developed during ripening, as shown in previous studies, this is the first study to determine the relationship between fruit parts and fruit characteristics. These results would provide fundamental evidence for the necessity of studies on the changes in fruit in differential zones.

